# The segregation of vocal circuits solves a credit assignment problem associated with multi-objective reinforcement learning

**DOI:** 10.1101/236273

**Authors:** Don Murdoch, Ruidong Chen, Jesse Goldberg

**Author notes:** **Author contributions:** DM carried out the experiments. DM, RC and JG designed the experiments and analyzed the data. JG wrote the paper. **Competing financial interests statement:** The authors declare no competing financial interests.

## Abstract

Motor circuits vary in topographic organization, ranging from a coarse relationship between neuron location and function to highly localized regions controlling specific behaviors. For unclear reasons, vocal learning circuits lie at this second extreme: they repeatedly evolved to be spatially segregated from other parts of the motor system. Here we show that spatially segregated motor circuits can solve a specific problem that arises when an animal tries to learn two things at once. We trained songbirds in vocal and place learning paradigms with brief strobe light flashes and noise bursts. Strobe light negatively reinforced place learning but did not affect song syllable learning. Noise bursts positively reinforced place preference but negatively reinforced syllable learning. These double dissociations indicate that vocalization-related reinforcement signals specifically target the vocal motor system, while place-related reinforcement signals specifically target the navigation system. Non-global, target-specific reinforcement signals have established utility in machine implementation of multi-objective learning. In vocal learners, such signals could enable an animal to practice vocalizing as it does other things such as forage for food or learn to walk.

## Introduction

Diverse behaviors can be shaped by primary reinforcement such as reward (e.g. food or water) and punishment (e.g. electric shock), including place preference, lever pressing, action sequencing and timing, reaching, choice tasks, and more^1^. Electrical or optogenetic activation of ascending neuromodulators such as dopamine can also reinforce a wide range of actions coincident with the stimulation^2,3^. The diffuse, non-topographic projection patterns of ascending neuromodulatory systems are well-suited to carry reinforcement signals globally to multiple action-generating modules in basal ganglia and cortex^4-6^.

Yet one problem with global reinforcement signals is credit assignment: how does the brain ‘know’ which action caused a reward and, relatedly, which action-generating neural circuit requires synaptic plasticity and policy updating to improve performance? Superstitious behaviors acquired during reinforcement learning exemplify how a global reinforcement signal can mis-assign credit to a motor act temporally contiguous with, but causally unrelated to reinforcement^7^. Stereotypic body rotations, arm and leg movements acquired during simple tapping or pecking tasks further demonstrate that motor regions controlling arm, leg, and orientation circuits share common, broadcasted reinforcement signals^8,9^.

The credit assignment problem is particularly severe in cases when an agent pursues multiple objectives at once^10-13^. For example, consider a toddler babbling to herself while stacking blocks. She uses her vocal motor system to speak and her hands and arms to stack. Learning these tasks depends on different types of feedback. Learning to talk may rely on comparison of sensory feedback to an internal auditory target, while learning to stack blocks may rely on comparison of sensory feedback to an entirely independent visual target.

Machine learning provides potential insights into reinforcement learning (RL)^14-16^. Whereas standard reinforcement learning (RL) algorithms optimize a single cost function (e.g. maximize cumulative reward) with a scalar reinforcement signal^17^, in multiobjective learning a single agent can be endowed with independent sub-agents which are trained by an equal number of agent-specific reinforcement signals^14-16^. In the babbling toddler, for example, auditory error signals would reach the vocal motor system (and not the block building one) to shape future vocalizations. Meanwhile errors such as tower collapse would reach the block-building system (and not the vocal motor one) to shape future block building policy^18^. To our knowledge it remains unknown if a single animal possesses distinct ‘agencies’ inside its brain which are, by definition, shaped by agent-specific reinforcement signals.

Here we use songbirds to test if an animal can compute behavior-specific reinforcement signals and route them to corresponding behavior-producing parts of the motor system. Songbirds sing and, at the same time, navigate (i.e. hop and fly). An objective of the song system is to produce a target sequence of sounds derived from the memory of a tutor song^19-21^. An objective of a navigation system is to avoid aversive stimuli^22^. Song learning can be reinforced with distorted auditory feedback (DAF): if a brief broadband sound is played to a bird as it sings a target syllable a certain way, the bird modifies its song to avoid the feedback^23,24^. A song-relevant reinforcement signal thus derives from auditory error^25-28^. Navigation policy can be reinforced with a bright strobe light: if a strobe is flashed in a specific place, many animals learn to avoid that place^29^. A navigation-relevant reinforcement signal can thus derive from an aversive visual stimulus.

Songbirds also have a discrete vocal motor ‘song system’, dedicated to song learning and production, that is spatially segregated from other parts of the motor system^30^. Lesions to song system nuclei impair singing but not other behaviors such as grooming, eating, navigation and flight^30-33^. In addition, neural activity in song system nuclei is strongly correlated with singing and not other motor behaviors^34-37^. Hummingbirds and parrots independently evolved the vocal learning capacity; curiously, they possess spatially segregated song systems^38-40^.

The ability of songbirds to simultaneously generate distinct behaviors with distinct objectives, together with the existence of a spatially isolated song system, presents a unique opportunity to test different network architectures for multi-objective learning. To determine if vocal and place learning can be shaped by shared, overlapping, or distinct reinforcers, we built a closed-loop system that provides either strobe light or noise feedback contingent on zebra finch spatial position or pitch of a target song syllable (Figure 1). As shown in Figure 2, distinct learning algorithms require distinct network architectures that make distinct and specific experimental predictions. In a standard RL network with a scalar, global reinforcement signal, both strobe and noise could similarly reinforce both song pattern and place preference (Figure 2A). In a multi-agent RL architecture where each behavior is independently trained by a behavior-specific reinforcement signal, noise could reinforce song pattern but not place preference, and strobe could reinforce place preference but not song pattern (Figure 2B). Finally, global and target-specific reinforcement signals might coexist: one of the stimuli could drive a global error signal that reinforces both behaviors, while another could specifically target one behavior (Figure 2C).

**Figure 1.**
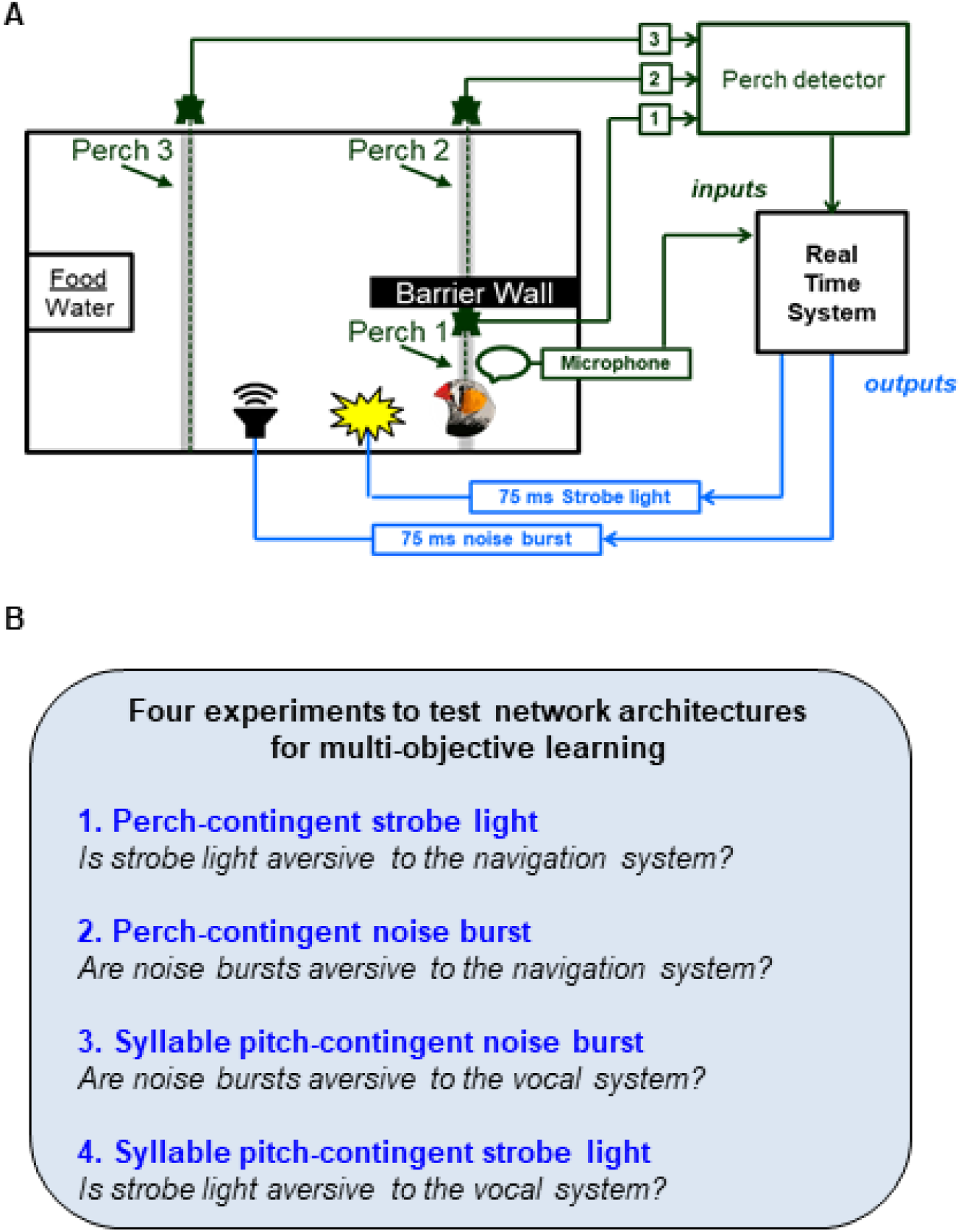
Experimental control of place preference and song syllable learning. **(A)** Schematic of experimental homecage. Signals from perch-mounted IR beam breaks and an overhead microphone provided inputs (green) to a system that analyzed perch occupancy and song features in real time. The system sent outputs (blue) to LEDs for strobe light feedback and to speakers for noise burst feedback. The system implemented one of four contingencies: (1) Perch contingent strobe light, to test if strobe influences place preference; (2) Perch contingent noise, to test of noise influences place preference; (3) Song syllable pitch contingent noise, to test if noise influences syllable selection; and (4) Syllable pitch contingent strobe, to test if strobe light influences syllable selection.

**Figure 2.**
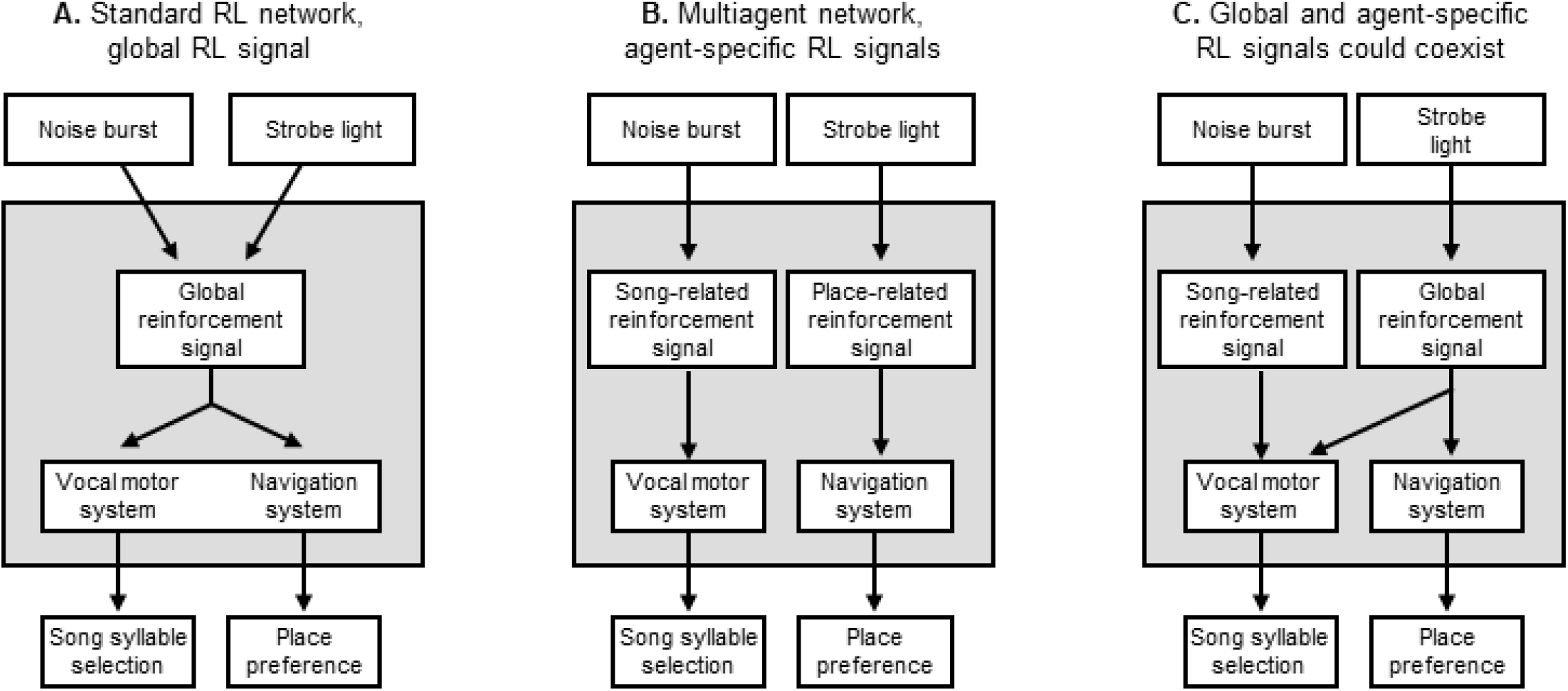
Different network architectures make specific predictions for how multiobjective reinforcement learning is implemented. **(A)** Schematic of a standard RL network where a single reinforcement signal acts globally on multiple parts of the motor system to shape the policy of multiple behaviors. This architecture predicts that both strobe light and noise burst will be aversive to both vocal motor and navigation systems, i.e. will shape both song syllables and place preference. **(B)** A multi-agent RL network where each behavior is shaped by its own behavior-specific reinforcement signal. This architecture predicts that noise will shape song but not place preference, and that strobe will shape place preference but not song. **(C)** Global and behavior-specific reinforcement signals might coexist. Here, it is imagined that strobe light drives reinforcement signals that reach all parts of the motor system, whereas DAF-related reinforcement signals target specifically the vocal motor system. This architecture predicts that DAF will shape song but not place preference, and that strobe will shape both song and place preference.

We find that song pattern and place preference are differentially reinforced by sound and strobe light respectively, consistent with a multi-agent network architecture. Our identification of behavior-specific reinforcement suggests that auditory feedback has privileged access to songbird vocal motor circuits. More generally, our results provide support for animal implementation of a specific network architecture used in machine learning and provide a logic for the spatial segregation of vocal motor circuits that independently evolved in diverse vocal learning species^16,41^.

## Results

To test if strobe light drives place learning, we implemented perch-contingent strobe light feedback: if a bird landed on one of two ‘target’ perches, a 75 millisecond strobe light stimulus discharged at 2 ± .25 Hz (see methods). Birds preferred to avoid the strobe-associated perch (Figure 3). Perch 1-contingent strobe resulted in preference for perch 2 (Perch 2 landing rate: 81.3±11.7%, p<0.0001; Perch 2 occupancy: 73.7±14.6%, p<0.01, one-sample t tests, n=6 birds). Contingency reversal with perch 2-contingent strobe biased preference towards perch 1 (Perch 1 landing rate increased from 18.7±11.7% to 59.4±21.4%, p<0.001; perch 1 occupancy increased from 26.3±14.6% to 73.7±14.6%, p<0.05; two-sample t-tests). These data indicate that strobe light negatively reinforce place preference.

**Figure 3.**
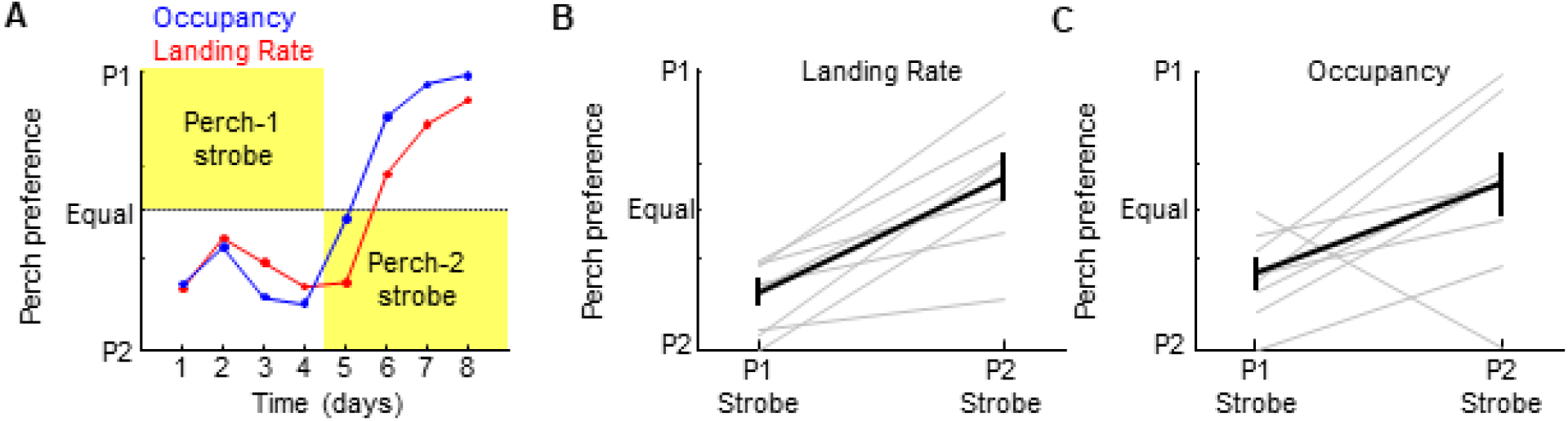
Strobe light is aversive to the navigation system. **(A)** Perch occupancy (blue) and landing rate (red) on test perches from an example bird, plotted over four days of perch 1-contingent strobe (P1 Strobe), followed by four days of perch 2-contingent strobe (P2 Strobe). **(B-C)** Average landing rates (B) and Occupancies (C) for six birds across P1- and P2-contingent strobe conditions demonstrate preference for non-strobed perch.

Perch-contingent auditory feedback was implemented exactly as described above except the 75 millisecond strobe was replaced with a 75 millisecond song-syllable like sound (Methods). Birds acquired a place preference for whichever perch triggered the noise (Figure 4). Perch 1-contingent noise resulted in preference for perch 1 (Perch 1 occupancy: 82.6 ± 14.4%, p<0.0001; Perch 1 landing rate: 76.0 ± 5.8%, p<0.01, one-sample t tests, n=6 birds). Contingency reversal with perch 2-contingent noise biased preference towards perch 2 (Perch 2 landing rate increased from 24.0±5.8% to 53.6±52.7%, p<0.05; perch 2 occupancy increased from 17.4±14.4% to 86.7±15.0%, p<0.001; two-sample t-tests). These data indicate that brief noise bursts positively reinforce place preference.

**Figure 4.**
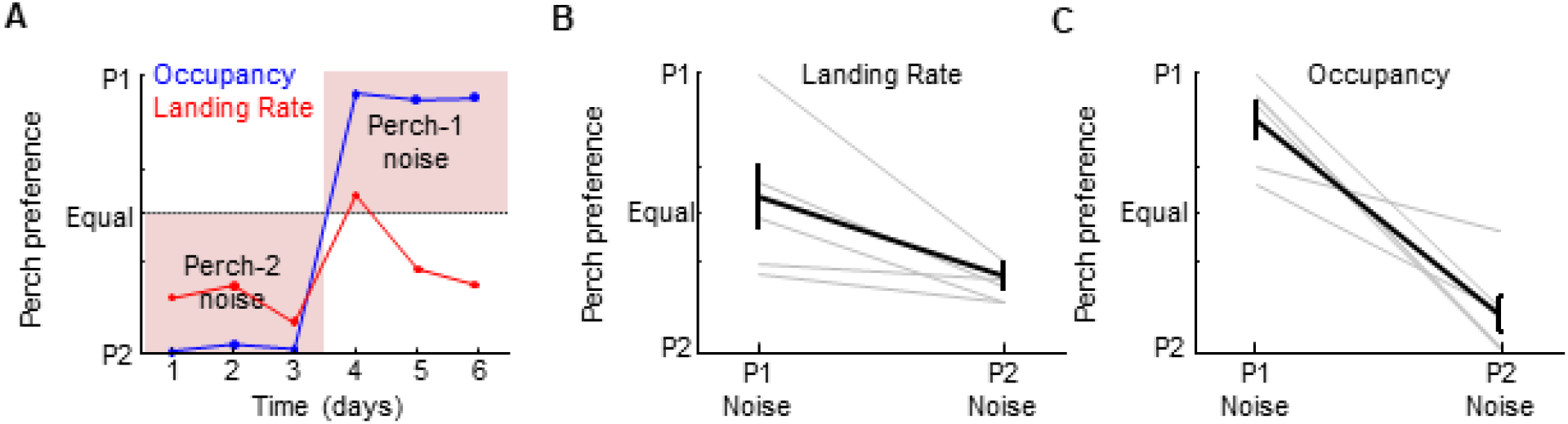
Noise bursts are positively reinforcing to the navigation system. **(A)** Perch occupancy (blue) and landing rate (red) on test perches from an example bird, plotted over four days of perch 2-contingent noise, followed by four days of perch 1-contingent noise. **(B-C)** Average landing rates (B) and Occupancies (C) for six birds across P1- and P2-contingent noise conditions demonstrate preference for the ‘noisy’ perch.

**Figure 5.**
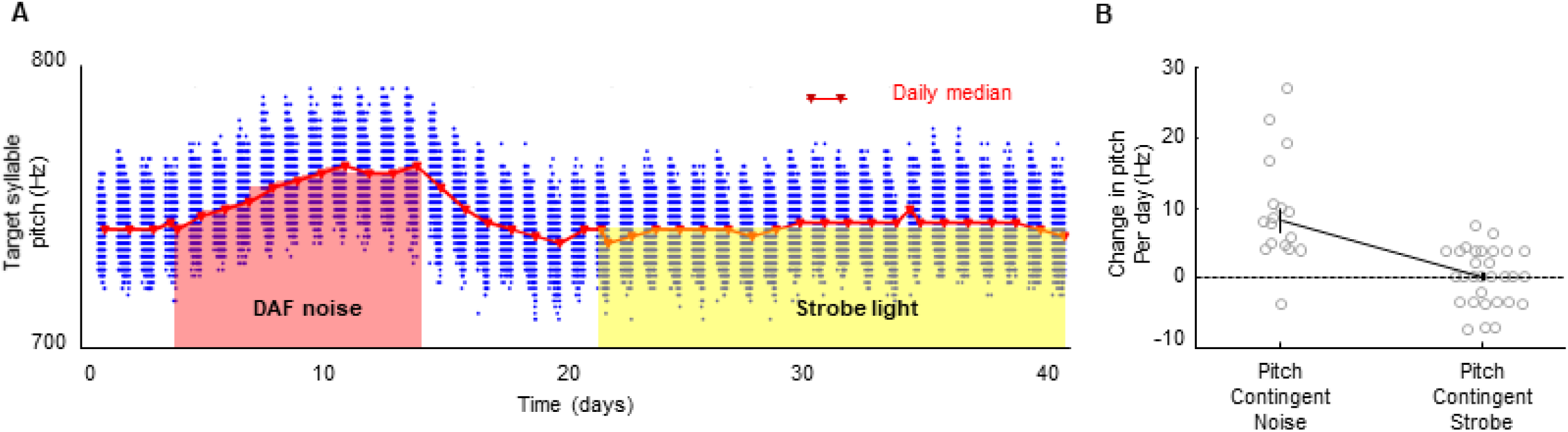
Noise feedback, but not strobe light, drives song syllable learning. **(A)** Blue dots denote mean pitch of target syllable renditions sung over 41 days for one bird. Pink and yellow shading demarcate syllable renditions that triggered noise and strobe light, respectively. **(B)** Change in mean pitch per day during pitch-contingent noise (left) or strobe light (right).

We next carried out song syllable pitch-contingent auditory feedback. In each bird, we chose a ‘target’ harmonic syllable amenable to real-time pitch computation (Methods). After at least three days of obtaining baseline target syllable pitch distributions, we implemented pitch-contingent noise feedback by playing the 75 ms noise burst (used in perch preference experiments) during low pitch target syllable variants (Figure 4). All birds increased the pitch of their target syllable to avoid the noise (change in pitch per day: 8.2±7.3 Hz, p<0.0001, one-sample t test, n=5 birds), consistent with previous studies^23,24,42-45^. Thus the same noise that was positively reinforcing to the navigation system was aversive to the vocal motor system.

To test if strobe light is aversive to the vocal motor system, we implemented pitch-contingent strobe feedback, exactly as described above except the 75 millisecond sound was replaced with the 75 millisecond strobe stimulus. Birds did not change the pitch of their target syllables to avoid strobe, even when they were given extended periods of time to allow for potentially slower learning (change in pitch per day: 0.2 ± 3.3 Hz, p>0.7, n = 5 birds, 45 days, one-sampled t test). Thus, the light stimulus that was aversive to the navigation system was not detectably aversive to the song system.

The routing of error signals to distinct parts of the motor system could in principle be gated by behavioral context. For example, the noise sound could be aversive during singing but not during non-singing periods (Figure 6A). To test this, we separately analyzed perch occupancy patterns for singing and non-singing periods during the perch-contingent noise experiments. Birds preferred the ‘noisy’ perch during both singing and non-singing periods (Figure 6B-D) (Two-way ANOVA showed significant effect of noise on perch occupancy [F(1,24)=114.65, p<0.000000001], and no effect of singing state [F(1,24)=0.21,p>0.6] or interaction between noise and singing state [F(1,24)=1.66, p>0.2]). Thus context-dependent gating of noise aversiveness cannot explain birds’ preference for occupying ‘noisy’ perches.

**Figure 6.**
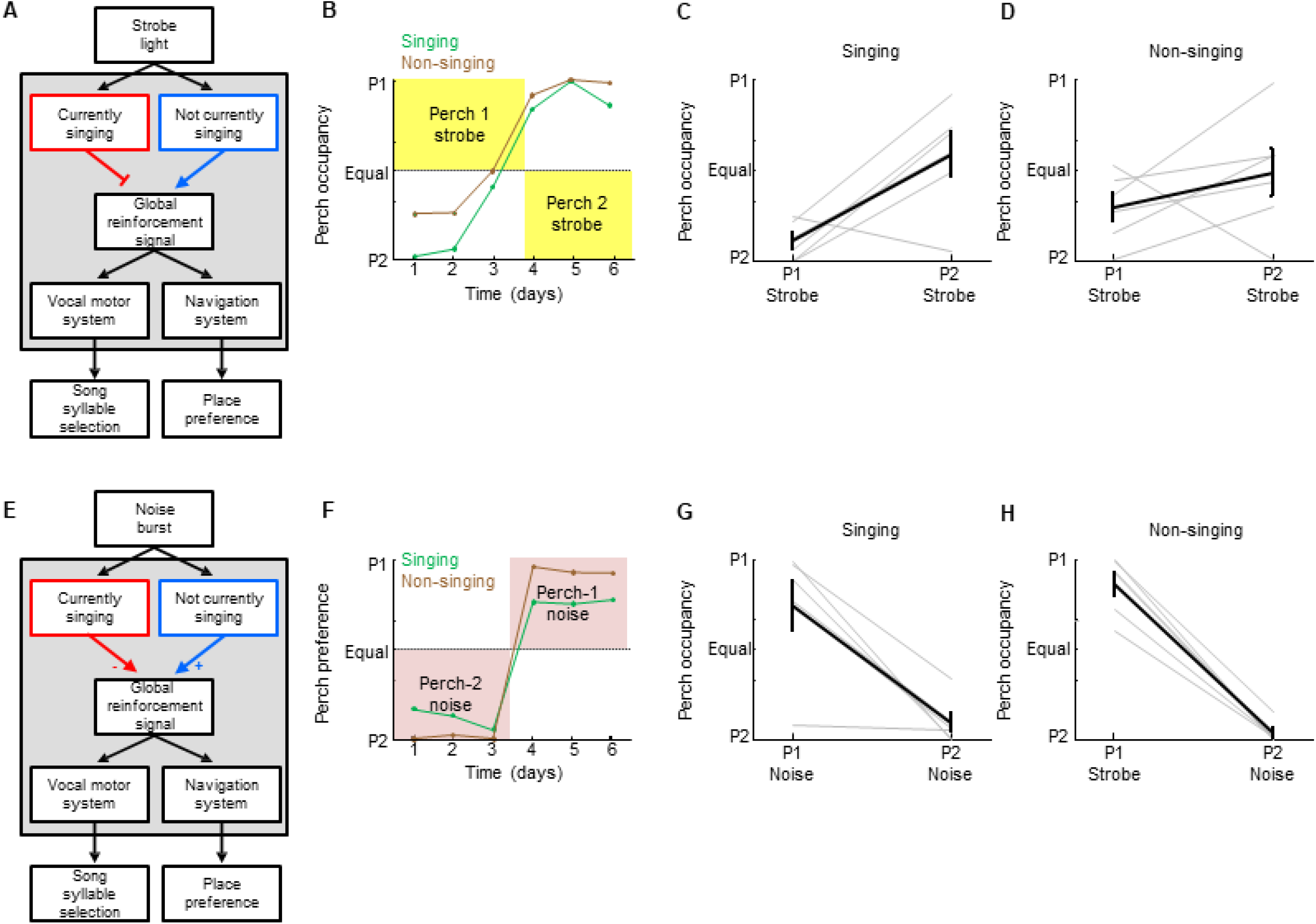
The reinforcing properties of noise and light do not depend on behavioral context. **(A)** A network architecture in which the access of strobe light to a global reinforcement signal is gated by singing state. This architecture would predict that strobe is not aversive when birds are singing. **(B)** Perch occupancy during singing (green) and non-singing (brown) on test perches from an example bird, plotted over three days of perch 1-contingent strobe (P1 Strobe), followed by three days of perch 2-contingent strobe (P2 Strobe). **(C-D)** Average perch occupancies during singing (C) and non-singing (D) for six birds across P1- and P2-contingent strobe conditions demonstrate preference for non-strobed perch during both singing and non-singing periods. **(E)** A network architecture in which the access of noise burst to a global reinforcement signal is gated by singing state. This architecture predicts that noise valence becomes negative during singing such that birds would not choose to sing on ‘noisy’ perches. **(F)** Perch occupancy during singing (green) and non-singing (brown) on test perches from an example bird, plotted over three days of perch 2-contingent noise, followed by three days of perch 1-contingent noise. **(G-H)** Average perch occupancies during singing (G) and non-singing (H) for six birds across P1- and P2-contingent noise conditions demonstrate preference for the noisy perch during both singing and nonsinging periods.

**Figure 7.**
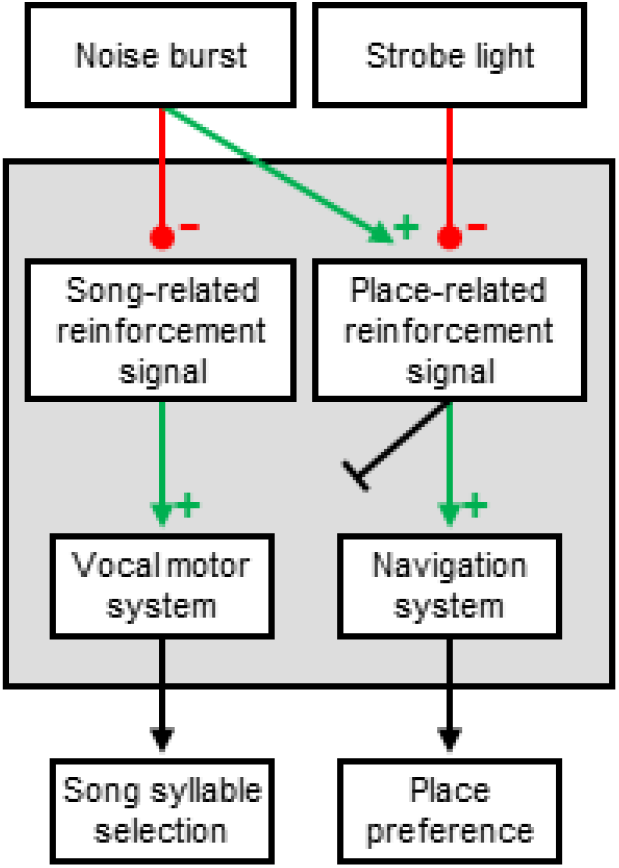
Network architecture supported by our experimental results. Noise burst was aversive to the vocal motor system and was reinforcing to the navigation system. Strobe light was aversive to the navigation system but was apparently unable to access vocal motor circuits.

Similarly, the strobe light might be globally aversive but only during non-singing periods, for example if birds simply did attend to light during singing (Figure 6E). To test this, we separately analyzed perch occupancy patterns for singing and non-singing periods during the perch contingent strobe experiments. Birds avoided the strobed perch during both singing and non-singing (Figure 6F-H) (Two-way ANOVA showed significant effect of strobe on perch occupancy [F(1,24)=15.26, p<0.001], and no effect of singing state [F(1,24)=0.23,p>0.6] or interaction between strobe and singing state [F(1,24)=2.82, p>0.1]). Thus context-dependent gating of strobe aversiveness cannot explain place preference for non-strobed perches.

## Discussion

Vocal learning poses unique problems because vocalizations are often produced as animals are doing other things. Toddlers babble even as they learn to walk; birds learn to sing even as they hop and fly around an environment. In machines, one way to solve the credit assignment problem associated with multi-objective reinforcement learning is to endow an agent with independent sub-agents which are trained by an equal number of agent-specific reinforcement signals^14-16^. In this view, functionally segregated vocal learning circuits could provide a target for vocalization-specific reinforcement that would not contaminate non-vocal behaviors. We report that song and place learning are driven by distinct reinforcers, demonstrating that action-specific reinforcement signals can be computed and precisely routed to corresponding sub-parts of the motor system. These findings also provide a clear counterexample to general purpose models of learning that rely on global reinforcement^4,46^.

Specific evolutionary histories endow animals with genetic constraints on the associativity of actions with outcomes^47^. For example, dogs struggle to learn to yawn for food, trapped cats readily learn to escape a cage by pressing a lever but not by grooming, rats associate sounds and lights with electric shock but not with nauseating food, and pigeons can learn to peck a key for food and take flight to avoid a shock, but not vice versa^1,48-50^. These studies demonstrate that pairing of specific actions with valent consequences in a laboratory setting may be so unnatural that an animal is unable, or ‘contraprepared’, to associate them^51^. In our experiments, it was likely natural for bird to navigate away from a threatening stimulus, but not to avoid eliciting it by singing in a different way. Reinforcing vocalizations based on auditory, but not visual feedback, may also be more natural for song imitation. Finally, it may also be natural for a social animal like a zebra finch to navigate towards noisy places and away from quiet ones, as silence may indicate isolation and an associated increased predation risk.

What are the precise neural circuits that connect an aversive light flash to the navigation system to drive avoidance behavior, and a song-like noise to the vocal motor system to change syllable pitch? First, much like the human speech system, the song system is a discrete neural circuit, embedded in an evolutionarily conserved basal ganglia thalamocortical loop^41,52^. Electrophysiology, brain lesion and immediate early gene studies indicate that the song system is dedicated to singing, and not to other behaviors such as grooming, eating or navigation^53^. The anatomical segregation of vocal circuits might create a discrete spatial target for song-specific error signals. For example, we recently identified song-related auditory error signals in dopaminergic neurons of the songbird ventral tegmental area (VTA)^25^. Using antidromic and anatomical methods we discovered that only a tiny fraction (<15%) of VTA dopamine neurons project to the vocal motor system - yet these were the ones that encoded vocal reinforcement signals. The majority of VTA neurons which project to other parts of the motor system did not encode any aspect of song or singing-related error. This specific ‘song evaluation channel’ embedded inside the ascending mesostriatal dopamine system thus targets auditory performance error signals specifically to vocal motor, and not navigation, circuits.

## Methods

### Animals

Subjects were 11 adult male zebra finches singly housed in behavior boxes singing undirected song. All experiments were carried out in accordance with NIH guidelines and were approved by the Cornell Institutional Animal Care and Use Committee.

### Pitch-contingent, syllable-targeted distorted auditory feedback

In five birds singing undirected song, song was recorded with AT803 Omnidirectional Condenser Lavalier Microphones amplified through a MIDAS xl48 8-Channel Microphone Pre-Amp connected to a National Instruments 6341 data acquisition card at 40 kHz using custom LabVIEW Software running on a windows PC (Dell Optiplex 7040 MT). The distorted auditory feedback (DAF) was a 75 millisecond duration broadband sound bandpassed at 1.5-8kHz, the same spectral range of zebra finch song^24^. Sound feedback was supplied as 16 bit 44.1 kHz wave file snippets using the Digilent High Performance Analog Shield (Digilent Part #410-309) through Logitech S120 Desktop Speakers. The amplitude was measured with a decibel meter (CEM DT-85A) and maintained at 88dB, less than the average peak loudness of zebra finch song^54^. Specific syllables were targeted either by detecting a unique spectral feature in the previous syllable (using Butterworth band-pass filters) or by detecting a unique inter-onset interval (onset time of previous syllable to onset time of target syllable) using the sound amplitude as previously described. In both cases a delay ranging from 10-200 ms was applied between the detected song segment and the precise part of the harmonic stack targeted for pitch-contingent DAF. We first determined the baseline pitch of each bird’s target harmonic syllable by recording song without distortion for at least 5 days. The pitch measured by taking a fast Fourier transform of a six millisecond segment within a specified portion of the harmonic stack^42^. The median pitch of the target syllable during day 5 of the baseline period was used as the initial threshold for feedback. On the first day of pitch-contingent DAF (day 6) we distorted target syllable renditions with pitch lower than this threshold. The distortion began 0-2 ms after the 6 ms window used for pitch measurement. Thresholds were automatically updated every 400 renditions if the median pitch of the last 400 renditions was higher than the previous threshold. We continued this protocol for several days until the birds moved their pitch up by at least 40 Hz from baseline (‘up’ days).

### Pitch-contingent, syllable targeted strobe light feedback

After pitch contingent distorted auditory feedback was demonstrably effective in inducing pitch changes, birds were given a zero-feedback epoch of at least 10 days during which their pitch distributions returned to baseline, as previously reported. Then pitch contingent syllable targeted light feedback was conducted exactly as described above, targeting the same syllables in the same five birds, except instead of playing the 75 ms DAF sound a 75 ms strobe light stimulus was flashed. Light feedback was delivered via custom LED panels with 24 LED’s per panel, 2 panels mounted on either end of each perch in a sandwich configuration (35000mcd per LED, manufacturer part #: LED Optek OVLEW1CB9, digikey part # 365-1177-ND). The strobe was a 75 millisecond event consisting of 5 ms of light-on, 65 ms of cage lights off, followed by 5 ms of light-on.

### Perch contingent DAF or strobe feedback

Six birds were taken from the colony and placed isolated in the test cages for 6-8 days of perch contingent strobe feedback (3-4 days per perch). The same birds were returned to the colony for at least 1 week and returned to test cages for 6-8 days of perch contingent noise (3-4 days per perch). Each perch was equipped with two 5mm IR-beam break sensors (Adafruit, product ID: 2168). Beam-break data was acquired and analyzed alongside the microphone signal with an arduino and custom labview code that communicated with either a speaker or strobe lights. Depending on the contingency, a targeted perch was associated with light or noise feedback.

### Statistical analyses

Statistics were first performed with two-way ANOVAs to test for effect of condition (strobe or no strobe, noise or no noise) and singing state (singing and non-singing), followed up with post hoc one-sample t tests to test whether specific conditions differed from the null hypothesis that perches would be equally occupied and landed on.

